## Supplemental Figures for "Short-term molecular consequences of chromosome mis-segregation for genome stability"

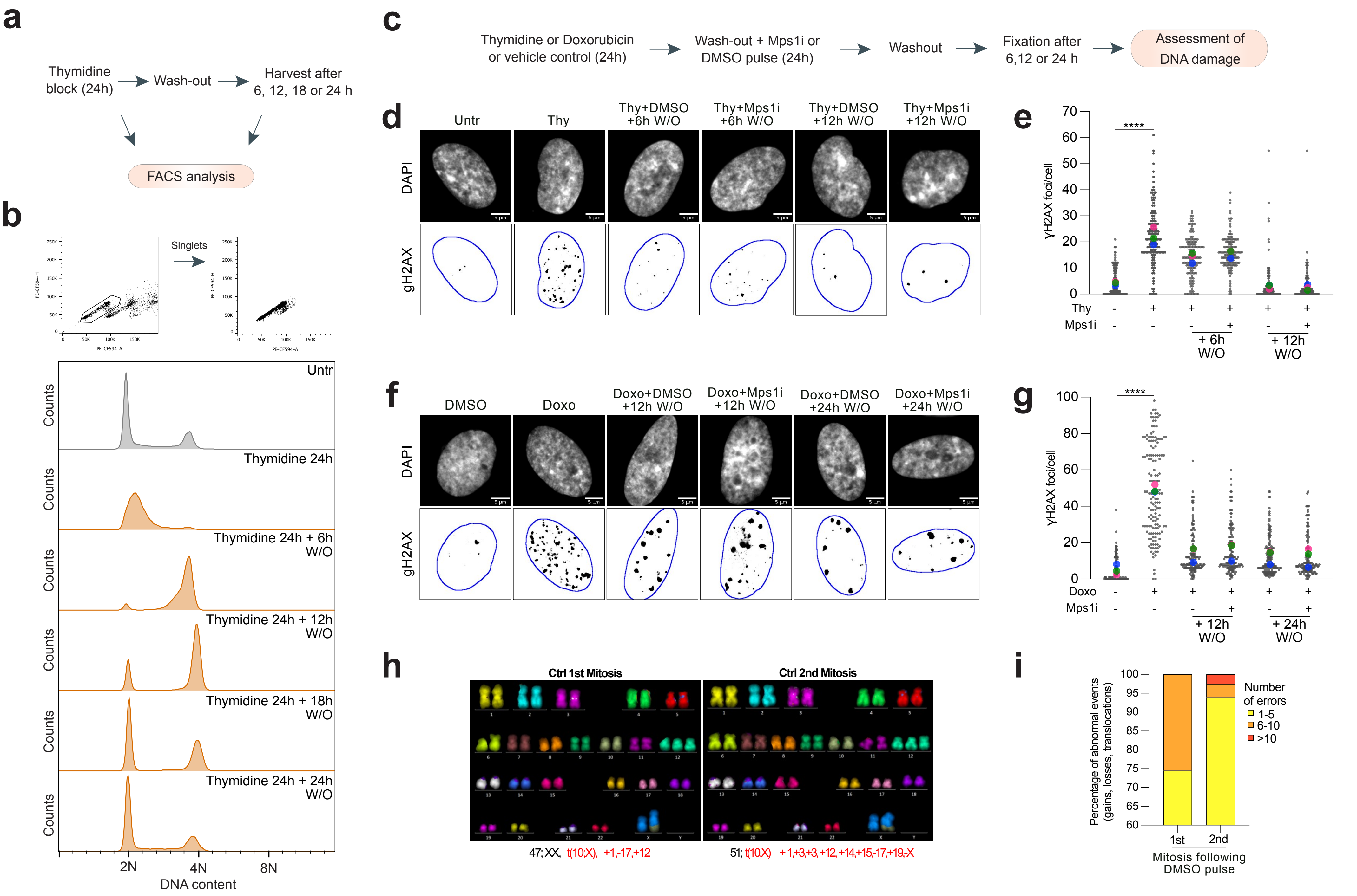


Extended Data Figure 1

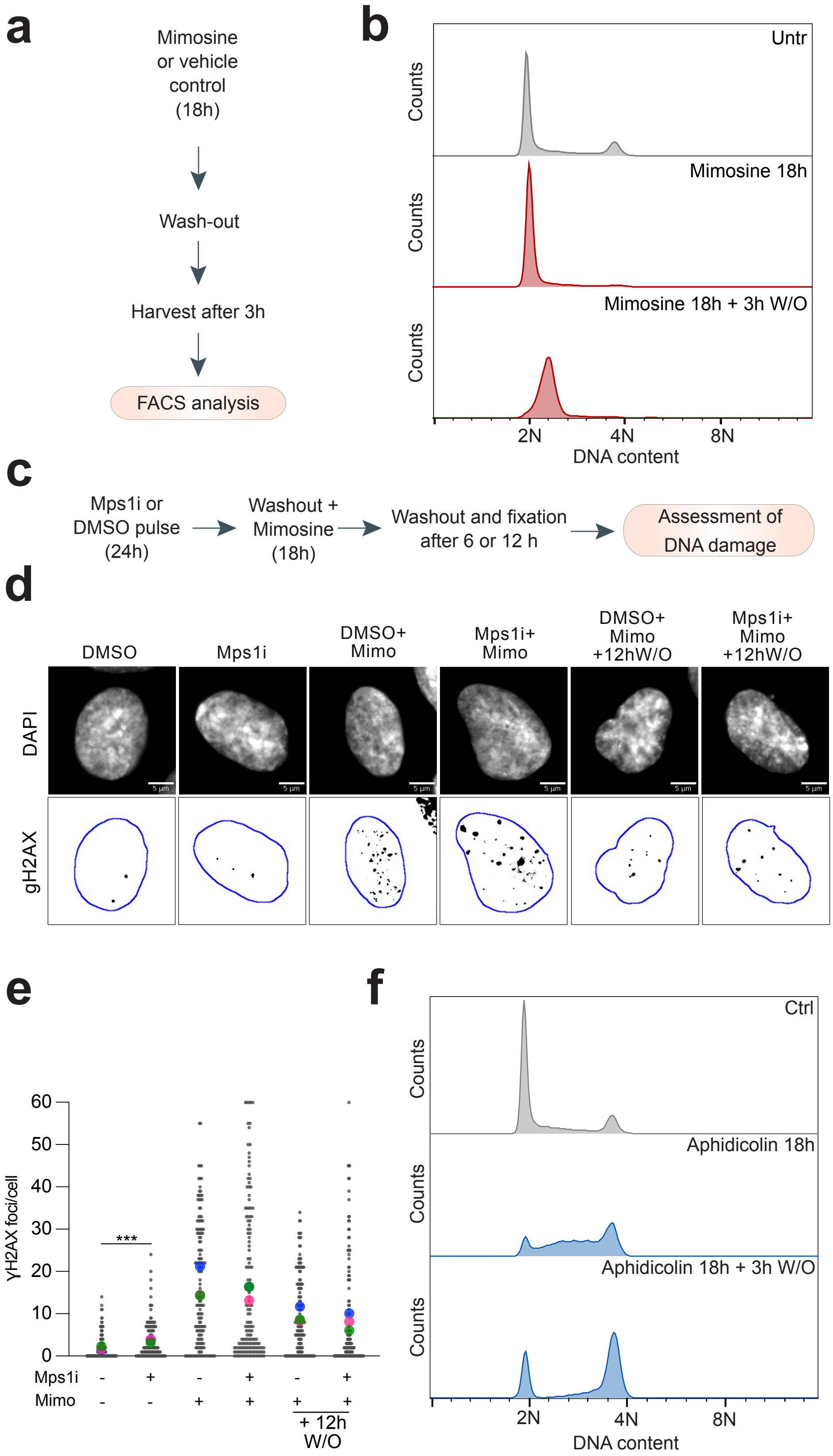


Extended Data Figure 2

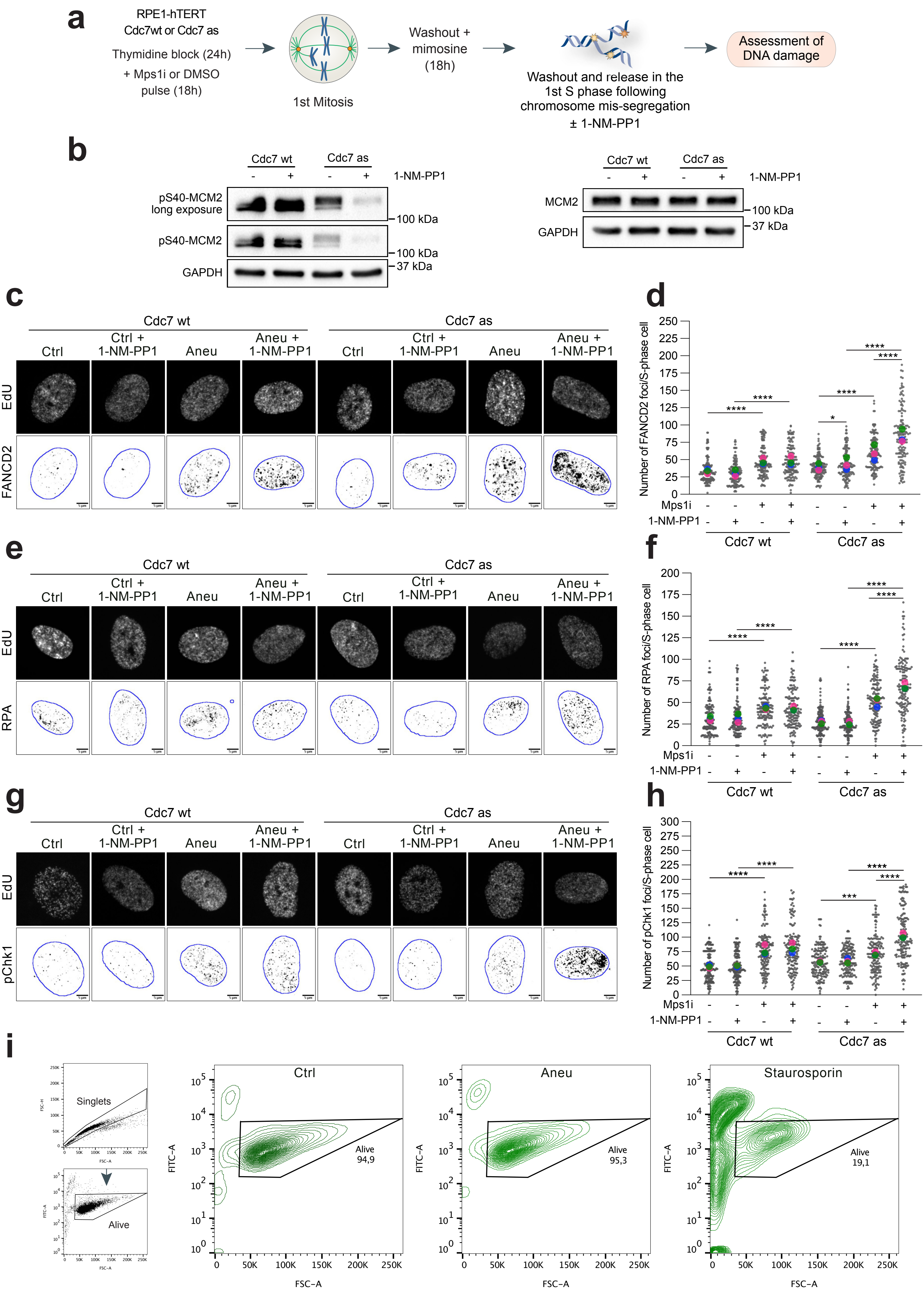


Extended Data Figure 3

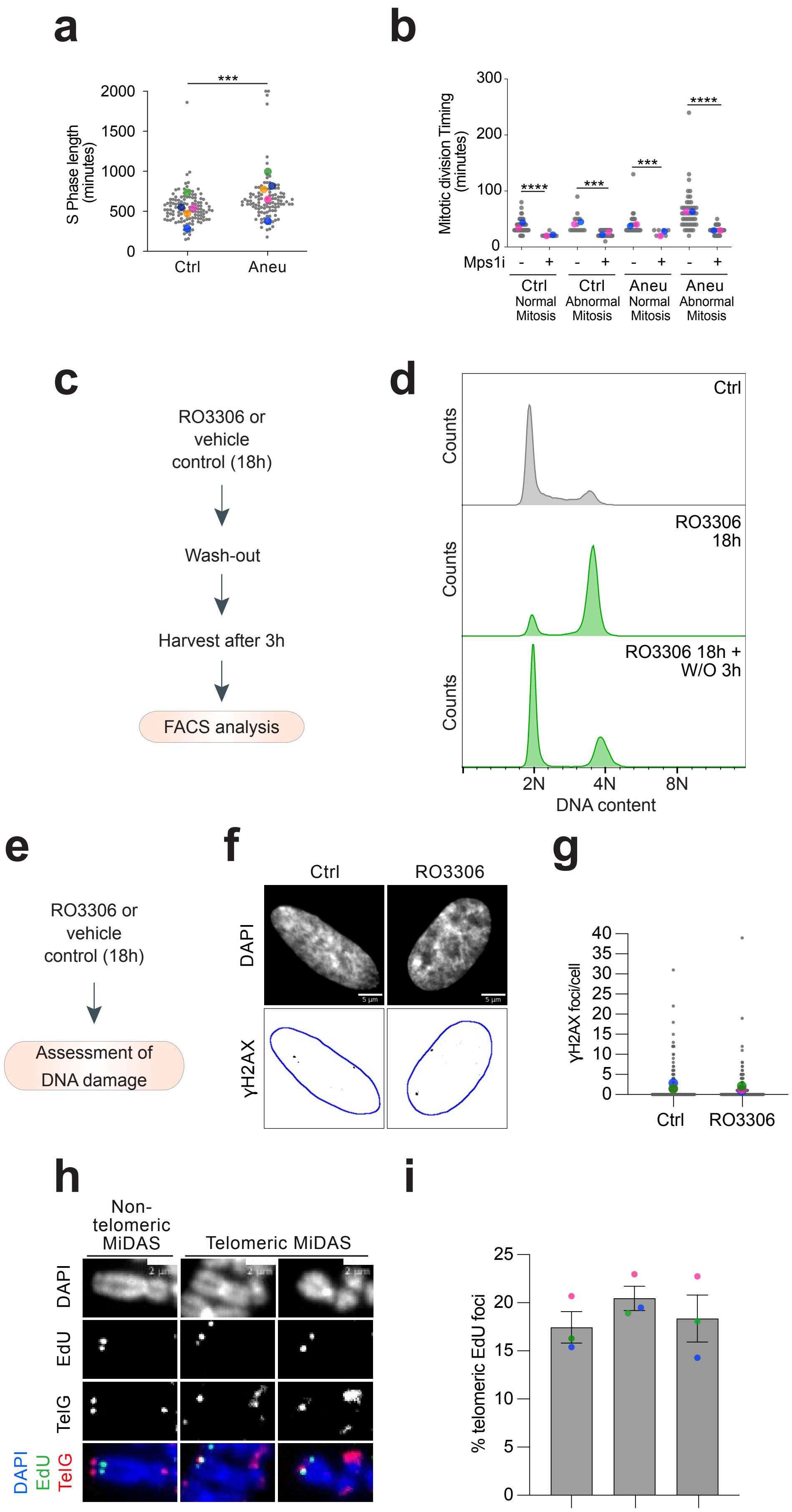


Extended Data Figure 4

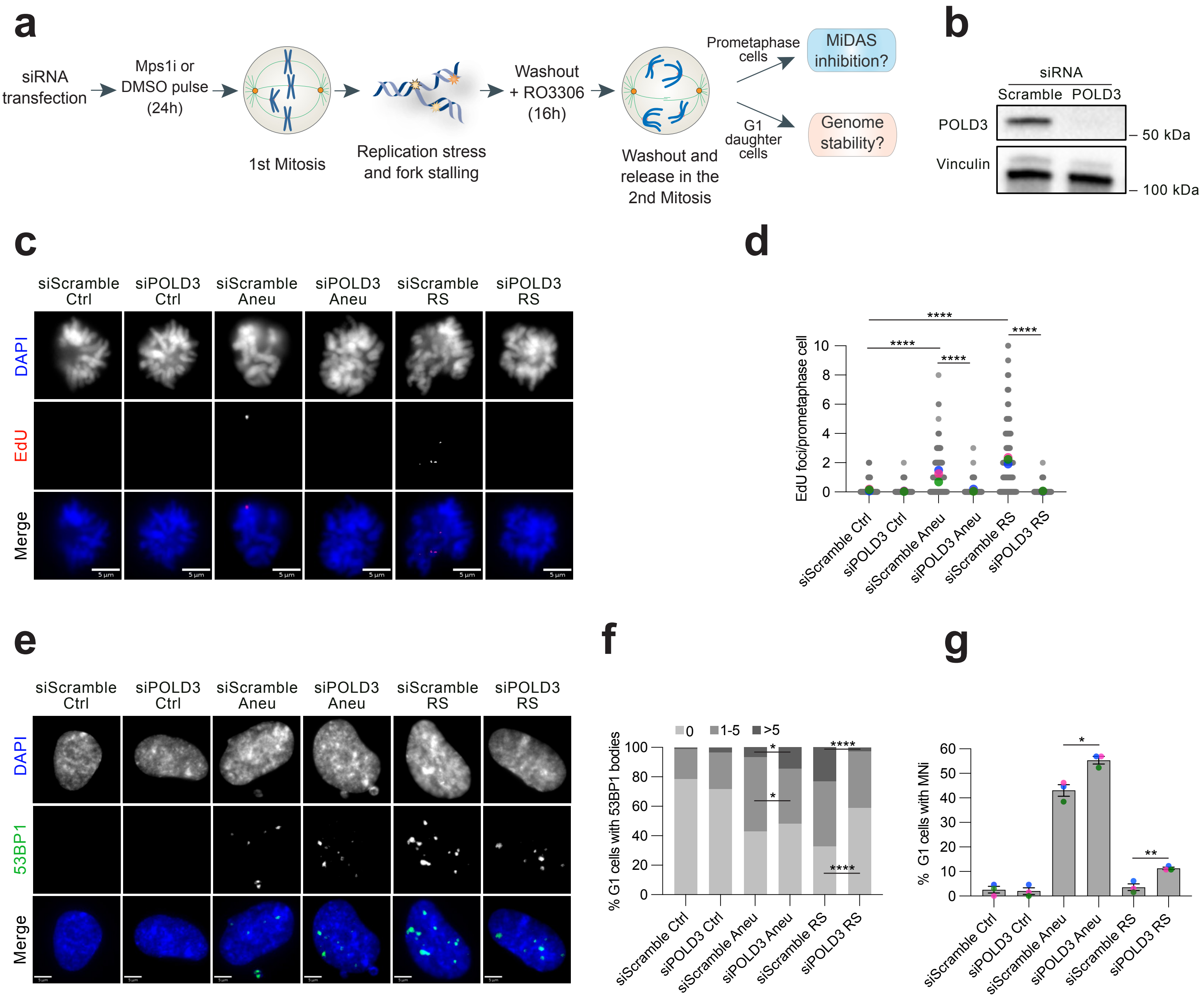


Extended Data Figure 5

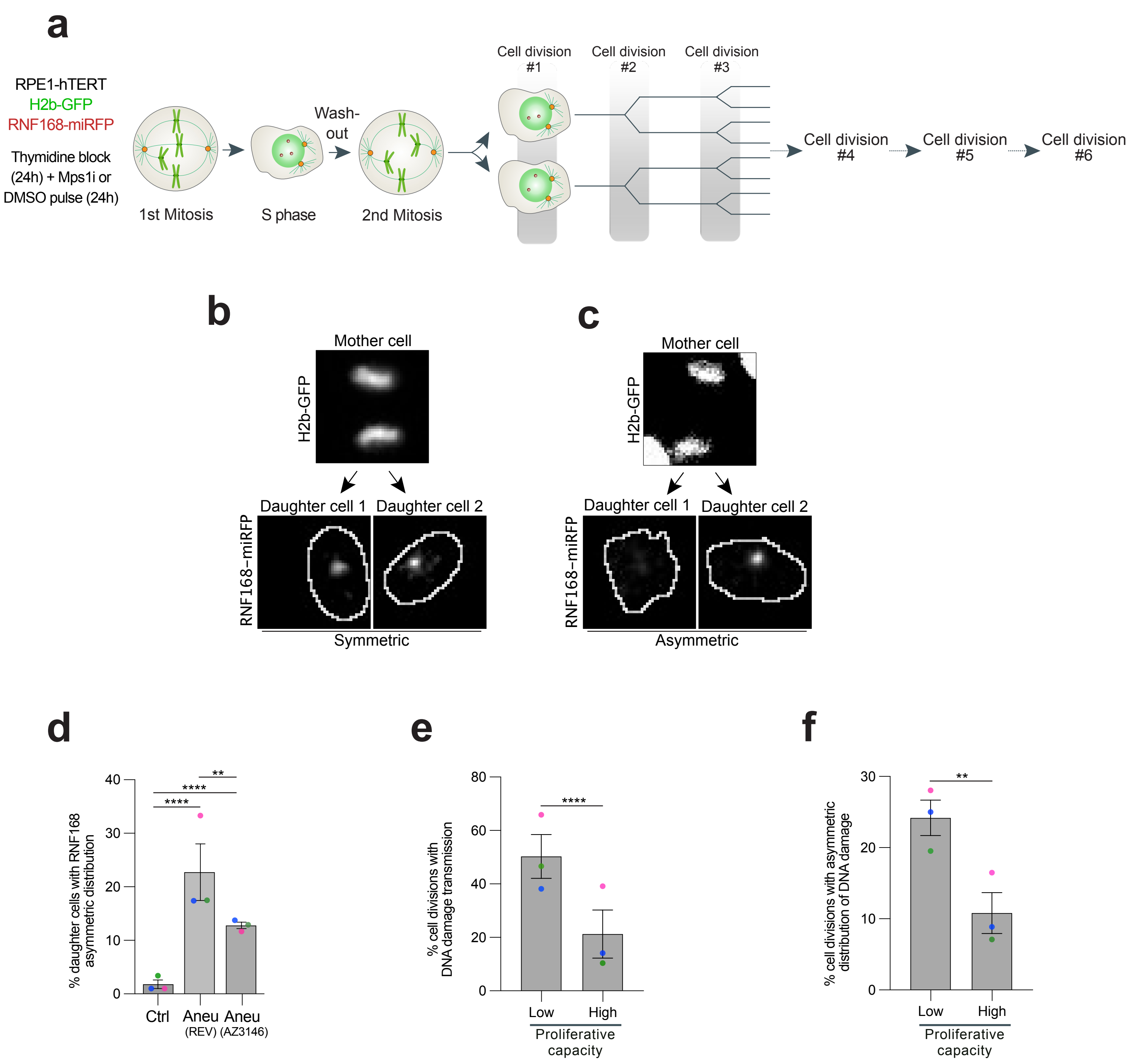


Extended Data Figure 6

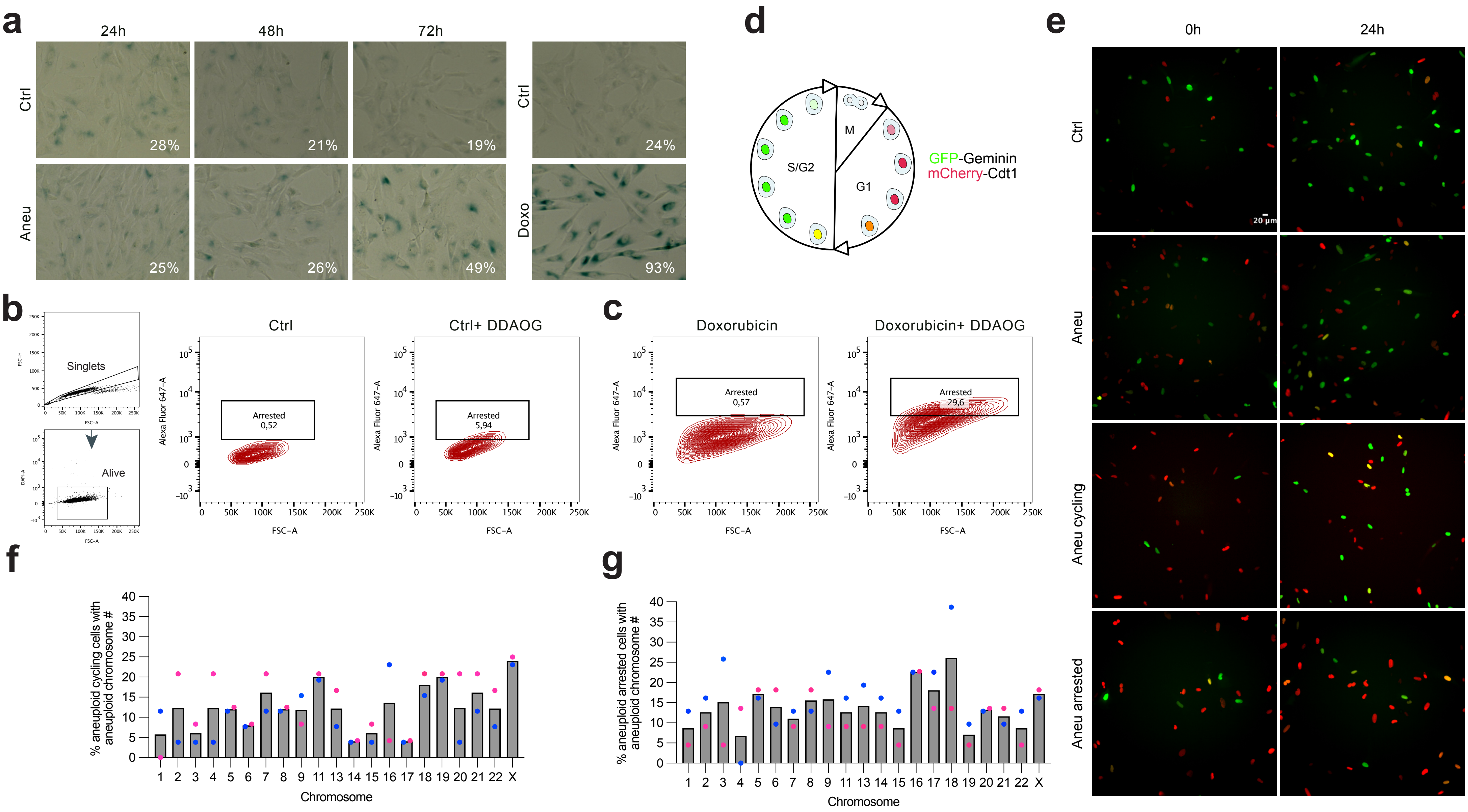


Extended Data Figure 7

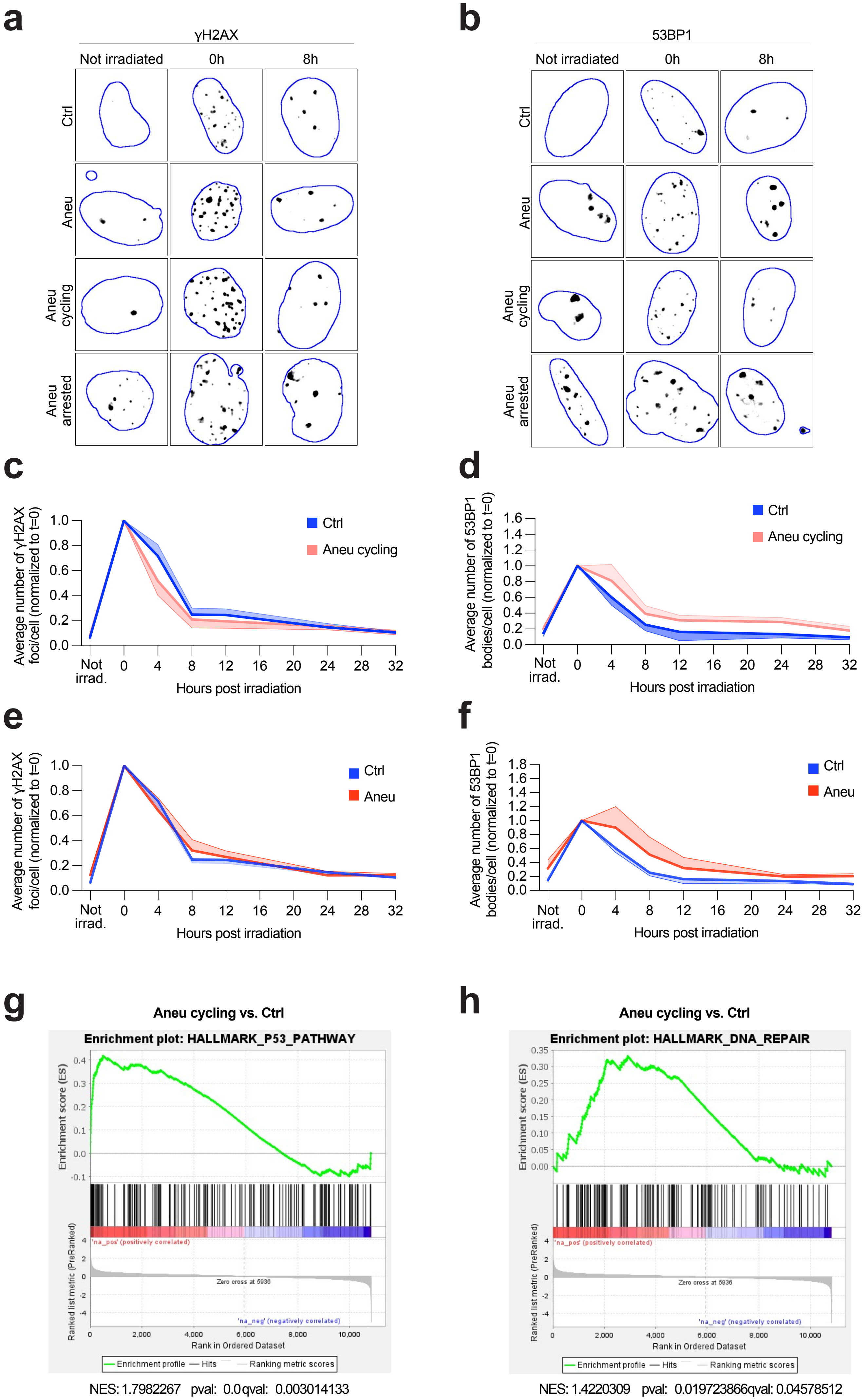


Extended Data Figure 8

| gene_name | ensembl_gene_id | baseMean | log2FoldChange | lfcSE | stat | pvalue | padj | signature |
|---|---|---|---|---|---|---|---|---|
| UHRF1 | ENSG00000276043 | 992,6407794 | -1,463494234 | 0,23690646 | -6,177519319 | 6,51166E-10 | 1,46332E-07 | GOBP_DNA_REPAIR [569] |
| FOXM1 | ENSG00000111206 | 1019,397201 | -1,365225147 | 0,581676165 | -2,347053616 | 0,018922532 | 0,137644121 | GOBP_DNA_REPAIR [569] |
| PRIM1 | ENSG00000198056 | 474,8477874 | -1,364119404 | 0,245844729 | -5,548703086 | 2,87796E-08 | 3,52553E-06 | HALLMARK_DNA_REPAIR [150] |
| FANCE | ENSG00000112039 | 267,7712184 | -1,343504755 | 0,283509612 | -4,738833176 | 2,14952E-06 | 0,000111839 | REACTOME_DNA_REPAIR [332] |
| H2BC14 | ENSG00000273703 | 174,2119743 | -1,330320082 | 0,388364543 | -3,425441649 | 0,000613801 | 0,010177337 | REACTOME_DNA_REPAIR [332] |
| PTTG1 | ENSG00000164611 | 2092,055002 | -1,311532721 | 0,195952716 | -6,693108159 | 2,1848E-11 | 9,23032E-09 | GOBP_DNA_REPAIR [569] |
| CCNA2 | ENSG00000145386 | 2017,460861 | -1,280223222 | 0,584722995 | -2,189452498 | 0,028563967 | 0,180154541 | REACTOME_DNA_REPAIR [332] |
| RAD54L | ENSG00000085999 | 95,30515172 | -1,264984213 | 0,41289 | -3,063731777 | 0,002185948 | 0,028823949 | KEGG_HOMOLOGOUS_RECOMBINATION [28] |
| RMI2 | ENSG00000175643 | 198,7630434 | -1,256125997 | 0,317193169 | -3,96012941 | 7,49092E-05 | 0,002013207 | REACTOME_DNA_REPAIR [332] |
| EME1 | ENSG00000154920 | 125,8010331 | -1,251873356 | 0,378354632 | -3,308730094 | 0,000937201 | 0,014493003 | KEGG_HOMOLOGOUS_RECOMBINATION [28] |
| HMGB2 | ENSG00000164104 | 2568,528678 | -1,247776619 | 0,623843451 | -2,000143812 | 0,045484737 | 0,242631208 | GOBP_DNA_REPAIR [569] |
| H4C1 | ENSG00000278637 | 384,8643662 | -1,239926814 | 0,314814741 | -3,938591977 | 8,19612E-05 | 0,002148074 | REACTOME_DNA_REPAIR [332] |
| BLM | ENSG00000197299 | 392,269428 | -1,238186534 | 0,264789399 | -4,676118227 | 2,92356E-06 | 0,000142957 | KEGG_HOMOLOGOUS_RECOMBINATION [28] |
| PIF1 | ENSG00000140451 | 89,12425074 | -1,225055521 | 0,430923506 | -2,842860752 | 0,00447106 | 0,049708774 | GOBP_DNA_REPAIR [569] |
| H2AX | ENSG00000188486 | 1633,516624 | -1,214070301 | 0,226237375 | -5,366356033 | 8,03433E-08 | 7,50961E-06 | REACTOME_DNA_REPAIR [332] |
| MCM2 | ENSG00000073111 | 1471,107745 | -1,212075792 | 0,212945245 | -5,691959872 | 1,25589E-08 | 1,72269E-06 | GOBP_DNA_REPAIR [569] |
| PCLAF | ENSG00000166803 | 400,8196156 | -1,211648027 | 0,262104055 | -4,622774828 | 3,78641E-06 | 0,000178536 | REACTOME_DNA_REPAIR [332] |
| GINS2 | ENSG00000131153 | 692,4481938 | -1,198244536 | 0,216352556 | -5,538388641 | 3,05267E-08 | 3,62273E-06 | GOBP_DNA_REPAIR [569] |
| CHAF1B | ENSG00000159259 | 475,6941015 | -1,192518656 | 0,27551539 | -4,328319574 | 1,50251E-05 | 0,000547535 | GOBP_DNA_REPAIR [569] |
| MCM7 | ENSG00000166508 | 3970,843886 | -1,192254667 | 0,206152086 | -5,78337426 | 7,32169E-09 | 1,14107E-06 | GOBP_DNA_REPAIR [569] |
| MCM3 | ENSG00000112118 | 2961,430599 | -1,179418929 | 0,179290935 | -6,57824071 | 4,76047E-11 | 1,7338E-08 | GOBP_DNA_REPAIR [569] |
| POLE2 | ENSG00000100479 | 114,5677156 | -1,155770914 | 0,393054504 | -2,940485102 | 0,003276988 | 0,039064947 | KEGG_BASE_EXCISION_REPAIR [35] |
| H2BC7 | ENSG00000277224 | 244,6185528 | -1,153603703 | 0,356981821 | -3,231547476 | 0,001231219 | 0,018061294 | REACTOME_DNA_REPAIR [332] |
| FANCA | ENSG00000187741 | 276,1320378 | -1,147377048 | 0,323144533 | -3,550662107 | 0,000384263 | 0,007145405 | REACTOME_DNA_REPAIR [332] |
| ESCO2 | ENSG00000171320 | 604,1535975 | -1,131052097 | 0,254374819 | -4,446399614 | 8,73214E-06 | 0,000354727 | GOBP_DNA_REPAIR [569] |
| CHAF1A | ENSG00000167670 | 652,2068713 | -1,126431655 | 0,225719477 | -4,990405206 | 6,02528E-07 | 3,97744E-05 | GOBP_DNA_REPAIR [569] |
| MCM4 | ENSG00000104738 | 3502,112152 | -1,118147714 | 0,178359277 | -6,269075182 | 3,63199E-10 | 9,35635E-08 | GOBP_DNA_REPAIR [569] |
| HMGB1 | ENSG00000189403 | 4022,455871 | -1,097626305 | 0,161937686 | -6,778078246 | 1,21785E-11 | 5,59257E-09 | KEGG_BASE_EXCISION_REPAIR [35] |
| BRCA2 | ENSG00000139618 | 475,1108442 | -1,095058696 | 0,230034854 | -4,760403373 | 1,93206E-06 | 0,000103063 | KEGG_HOMOLOGOUS_RECOMBINATION [28] |
| DTL | ENSG00000143476 | 1403,11263 | -1,074765995 | 0,192323372 | -5,588327526 | 2,29267E-08 | 2,98953E-06 | REACTOME_DNA_REPAIR [332] |
| MCM5 | ENSG00000100297 | 1618,342543 | -1,072958858 | 0,202528225 | -5,297823822 | 1,17191E-07 | 1,01457E-05 | GOBP_DNA_REPAIR [569] |
| MCM6 | ENSG00000076003 | 1350,168882 | -1,070443972 | 0,191730505 | -5,583065521 | 2,36316E-08 | 3,04386E-06 | GOBP_DNA_REPAIR [569] |
| XRCC3 | ENSG00000126215 | 153,407276 | -1,067855756 | 0,409861183 | -2,605408369 | 0,009176482 | 0,083650383 | KEGG_HOMOLOGOUS_RECOMBINATION [28] |
| PAXX | ENSG00000148362 | 273,4354076 | -1,066297861 | 0,285542718 | -3,734284907 | 0,000188249 | 0,004057733 | GOBP_DNA_REPAIR [569] |
| ZWINT | ENSG00000122952 | 1208,569743 | -1,057468524 | 0,216065679 | -4,894199437 | 9,87067E-07 | 6,06128E-05 | HALLMARK_DNA_REPAIR [150] |
| H2BU1 | ENSG00000196890 | 164,4035717 | -1,0526433 | 0,425867637 | -2,471761665 | 0,013444912 | 0,107579665 | REACTOME_DNA_REPAIR [332] |
| DDX11 | ENSG00000013573 | 451,5202913 | -1,051661549 | 0,247909922 | -4,24211158 | 2,21427E-05 | 0,000754422 | GOBP_DNA_REPAIR [569] |
| H2AZ1 | ENSG00000164032 | 7719,34297 | -1,050492077 | 0,167503262 | -6,271472355 | 3,5765E-10 | 9,35635E-08 | REACTOME_DNA_REPAIR [332] |
| TRIP13 | ENSG00000071539 | 869,8179167 | -1,046154634 | 0,226576705 | -4,617220622 | 3,88914E-06 | 0,000182389 | GOBP_DNA_REPAIR [569] |
| RAD51 | ENSG00000051180 | 257,3393749 | -1,029007726 | 0,300546848 | -3,423784787 | 0,000617555 | 0,010223537 | HALLMARK_DNA_REPAIR [150] |
| DNA2 | ENSG00000138346 | 657,8128943 | -1,018641876 | 0,215024678 | -4,737325439 | 2,16557E-06 | 0,000112121 | REACTOME_DNA_REPAIR [332] |
| UBE2T | ENSG00000077152 | 667,2240543 | -1,00051634 | 0,248650561 | -4,023784771 | 5,72703E-05 | 0,001617349 | REACTOME_DNA_REPAIR [332] |
| CLSPN | ENSG00000092853 | 757,2423379 | -0,999979178 | 0,221806732 | -4,508335558 | 6,53382E-06 | 0,000289959 | REACTOME_DNA_REPAIR [332] |
| FANCD2 | ENSG00000144554 | 846,2549826 | -0,990211718 | 0,201856287 | -4,905528246 | 9,31762E-07 | 5,78898E-05 | REACTOME_DNA_REPAIR [332] |
| POLA1 | ENSG00000101868 | 675,0218176 | -0,984358017 | 0,20494672 | -4,802994741 | 1,5631E-06 | 8,6437E-05 | HALLMARK_DNA_REPAIR [150] |
| MCM8 | ENSG00000125885 | 296,7887492 | -0,977502892 | 0,267888176 | -3,648921385 | 0,000263344 | 0,005247992 | GOBP_DNA_REPAIR [569] |
| RMI1 | ENSG00000178966 | 149,590574 | -0,973469008 | 0,363808538 | -2,675772851 | 0,007455716 | 0,072645083 | REACTOME_DNA_REPAIR [332] |
| BRCA1 | ENSG00000012048 | 987,4511882 | -0,970000371 | 0,189075516 | -5,130227294 | 2,89393E-07 | 2,12261E-05 | REACTOME_DNA_REPAIR [332] |

Extended Data Table 1
